# Epigenomic and genomic landscape of *Drosophila melanogaster* heterochromatic genes

**DOI:** 10.1101/239483

**Authors:** Parna Saha, Divya Tej Sowpati, Rakesh K Mishra

## Abstract

Heterochromatin is associated with transcriptional repression. In contrast, several genes in the pericentromeric regions of *Drosophila melanogaster* are dependent on this heterochromatic environment for their expression. Heterochromatic genes encode proteins involved in various developmental processes. Several studies have shown that a variety of epigenetic modifications is associated with these genes. Here we present a comprehensive analysis of the epigenetic landscape of heterochromatic genes across all the developmental stages of *Drosophila* using the available histone modification and expression data from modENCODE. We find that heterochromatic genes exhibit combinations of active and inactive histone marks that correspond to their level of expression during development. Thus, we classified these genes into three groups based on the combinations of histone modifications present. We also looked for potential regulatory DNA sequence elements in the genomic neighborhood of these genes. Our results show that Nuclear **M**atrix **A**ssociated **R**egions (MARs) are prominently present in the intergenic regions of heterochromatic genes during embryonic stages suggesting their plausible role in pericentromeric genome organization. We also find that the intergenic sequences in the heterochromatic regions have binding sites for transcription factors known to modulate epigenetic status. Taken together, our meta-analysis of the various genomic datasets suggest that the epigenomic and genomic landscape of the heterochromatic genes are distinct from that of euchromatic genes. These features could be contributing to the unusual regulatory status of the heterochromatic genes as opposed to the surrounding heterochromatin, which is repressive in nature.

## Introduction

Eukaryotic genomes can be broadly classified into euchromatin and heterochromatin. While euchromatin is gene rich and transcriptionally active, heterochromatin is repeat-rich, gene scarce and refractory to transcription. Constitutive heterochromatin present at the centromeres and telomeres, is marked by the repressive histone modifications - H3K9me3 and H4K20me3 and Su(var) proteins like HP1a. Constitutive heterochromatin accounts for nearly 30% of the genome in *Drosophila melanogaster* [1] but has remained relatively less studied as it is believed to have fewer functional features. However there has been a paradigm shift in the recent years, as active transcription of protein coding genes from the constitutive heterochromatin has been reported in different species like fly, mouse, humans and plants [2–6]

Heterochromatic genes encode various proteins involved in different cellular functions like ribosomal proteins, kinases, transporter proteins [7]. *Drosophila* is an excellent model with robust genetic techniques that made studies on genes located in deep heterochromatin possible even before the advent of genomic technologies. Heterochromatic genes were initially discovered in *Drosophila melanogaster* while searching for complementation groups in Ethyl methanesulfonate mutagenesis screen and well-known heterochromatic genes – *light* and *rolled* were mapped on either side of the centromere on chromosome 2 [8]. Subsequently many other heterochromatic genes were characterized on the autosomal chromosomes [9–17]. Several heterochromatic genes were also identified on the sex chromosomes [18]. 4^th^ chromosome although small in size, has peculiar features with respect to the heterochromatin. While some parts of the 4^th^ chromosome is facultative heterochromatin, majority of it (~75%) is interspersed with constitutive heterochromatic patches and active genes between those domains [19]. With the improvements in genome-sequencing technologies and the Drosophila Heterochromatin Genome Project (DHGP) [20] the number of centromeric heterochromatic genes in *Drosophila melanogaster* is currently in hundreds [1]. Upon comparing the structure of *D melanogaster* heterochromatic genes with their orthologs in other species of *Drosophila* where they are located in the euchromatin, promoter sequences were unchanged. However, in *Drosophila melanogaster* these genes have larger introns due to integration of transposable elements.[21]. This indicates that as a measure to counter-act genomic invasions by transposable elements, these genes are now in the heterochromatin but have evolved newer regulatory mechanisms to retain the functional status. Such regulatory mechanisms utilizes the specific features of the genomic and epigenomic landscape of heterochromatin.

Interestingly, not only do heterochromatic genes reside in repressive domains but they also require the heterochromatic environment for transcription. Chromosomal rearrangements that translocated these genes into the euchromatin suppressed their expression levels [22–23]. These genes showed ‘reciprocal’ heterochromatic Position Effect Variegation (PEV), more prominently in the background of Su(var) mutants as in case of *rolled* and *light* [24]. It was also reported that combinations of Su(var) mutations – Su(var)205 (HP1a), Su(var)208 and Su(var)210 could reduce eye pigmentation due to lowering of *light* expression even in the absence of chromosomal rearrangements [25,26]. Expression of both *Rpl15* (essential gene) and *Dbp80* (non-essential genes) were compromised in the genetic background deficient for HP1a [14]. These experiments confirmed that the genes located in heterochromatin require the heterochromatic environment for their expression. However, the mechanisms of transcriptional regulation of heterochromatic genes are still unknown

In this study, we have explored the DNA sequence and epigenetic features at the heterochromatic genes of *Drosophila melanogaster* to understand how heterochromatin not only permits but also is essential for the transcription of genes located within it. Customizing our epigenomic analysis tool, C-State [27], we have created an interactive *Drosophila melanogaster* heterochromatic gene database - Drosophila Het C-State. It uses published whole genome datasets to provide a platform for integrating the histone mark trends at heterochromatic genes with their expression profiles at different developmental stages. We find that heterochromatic genes can be broadly classified into three groups based on the combinations of histone marks associated with them in their highest and lowest expression stages. We also determined the characteristics of nuclear-matrix associated regions (MARs) present in the pericentromeric heterochromatin. Furthermore, we investigated the intergenic sequences for enrichments of any transcription factor binding sites. These results provide novel insights into the various genomic and epigenomic features of *Drosophila* heterochromatic genes and sets the stage for further experimental investigations.

## 2 Materials and methods

### 2.1 Retreiving datasets for epigenomic meta-analysis

60 well characterised (annotated gene structure, function and expression profile known) pericentromeric heterochromatic genes which were chosen. Gene expression data for these genes, normalized for both sequencing depth and transcript size (RPKM values, Supplementary Table 1: Stagewise expression file), were retrieved from the Flybase Temporal RNA Expression Data [28] using DGET [29]. Use of RPKM values allowed us to compare the expression levels of genes across various developmental stages directly. For each gene, a gene-epicard (available at the Table Summary of the tool webpage) was created that contains information like location, size, expression values in each stage, expression pattern and functions. For comparison of dynamicity of the epigenetic landscape over each gene, the histone modification ChIP-seq (generated by Kevin White lab, University of Chicago as part of modENCODE project) data for all developmental stages were downloaded from modENCODE [30]. We used ChIP-chip data where ChIP-Seq data was unavailable for a given histone modification/developmental stage. To facilitate comparison across various studies, we used the interpreted data files that provide enrichment of various histone modifications as genome coordinates. They were scaled to same parameters for evaluation of enrichment or lack of peaks.

**Table1:** List of the datasets (with their modencode accession numbers) taken for epigenomic analysis using C-State.

| **Marks** | **H3K9me3** | **H3K27me3** | **H3K4me1** | **H3K4me3** | **H3K9ac** | **H3K27ac** |
| --- | --- | --- | --- | --- | --- | --- |
| **Stages** |  |  |  |  |  |  |
| **E0-4 hrs** | 801 | 810 | 777 | 789 | 821 | 834 |
| **E4-8 hrs** | 802 | 811 | 778 | 790 | 822 | 835 |
| **E8-12 hrs** | 803 | 812 | 779 | 791 | 823 | 836 |
| **E12-16 hrs** | 804 | 813 | 780 | 792 | 824 | 837 |
| **E16-20 hrs** | 805 | 814 | 781 | 793 | 825 | 838 |
| **E20-24 hrs** | 806 | 815 | 782 | 794 | 826 | 839 |
| **L1** | 807 | 816 | 783 | 795 | 827 | 840 |
| **L2** | 808 | 817 | 784 | 796 | 828 | 841 |
| **L3** | 358* | 818 | 785 | 797 | 829 | 842 |
| **PUPAE** | 809 | 819 | 786 | 798 | 830 | 843 |
| **FEMALE** | 429* | 346 | 787 | 799 | 831 | 844 |
| **MALE** | 425* | 820 | 788 | 800 | 832 | 845 |

\*ChIP on chip dataset

### 2.2 Analysis of matrix associated regions (MARs)

Raw data from our previously published embryo 0–16hrs MAR sequencing (NCBI SRA Accession number-SRX443533) was used.. Previously unmapped dm3 MARs were mapped to the heterochromatic regions of dm6 build (better annotation of pericentromeric arm heterochromatin) [31] using Flybase Coordinate converter [32] to allow for inclusion of most MARs falling in heterochromatin. Supplementary Table 2 shows the coordinates of pericentromeric region used in the study. The MARs (Supplementary Table 3: Het MARs) were compared to other genomic features like introns, exons and intergenic regions

**Table2:** Grouping of the heterochromatic genes based on their dependency on inactive mark H3K9me3 for expression.

| Groups | Characteristic | Examples |
| --- | --- | --- |
| I | Active and inactive marks in stages of highest expression.<br>(a) Only inactive mark in stages of lowest expression<br>(b) active and inactive marks in stages of lowest expression | (a) *Atf6*, **CG10395**, *CG10417*, *CG12547*, *CG12567*, *CG14464*, *CG17493*, *CG17528*, *CG17683*, *CG40191*, **CG40439**, *Fis1*, *IntS3*, *Nipped-B*, *Parp*, *Rpl15*, *Rpl38*, *Spf45*, *Stlk*, *alpha-cat*, *gus*, *light*, **sxc**, *uex*,<br>(b) *CG11665*, *CG3107*, *CG10465*, *CG17486*,, *casein kinaseII* *alpha*, *Rpl5*, *cta*, *d4*, *Gprk1*, *CG17691* |
| II | active marks present in stages of highest and inactive (or active in case of constitutively expressing genes) in stages of lowest expression | *CG30438*, **CG32457**, *Haspin*, *gek*, *rl*, *Dbp80* |
| III | Only inactive marks present in stages of both highest and lowest expression | **CG17684**, *CG17508*, *CG30440*, **Nipped-A**, *Cht3*, *p120ctn*, *vtd*, *CG40006*, **CG40498**, **CG41378**, *Gpb5*, **CG41520**, **Scp1**, **TpnC4** **TpnC41c**, *nrm*, *Snap25* |

\* CG17490, CG17715: expression data Not Available
\*\* Genes with **Tissue specific expression** marked in red, constitutively expressing genes in black

**Table3:** Comparison of the characteristics of euchromatic and pericentromeric Het MARs.

| Properties | Euchromatic MARs | Pericentromeric Het MARs |
| --- | --- | --- |
| Average size (in bp) | 600 | 354 |
| Average inter-MAR distance | 16 kb | 45 kb |
| Location | Intronic | Intergenic |
| Proximity to DNaseI HSS site | NA | 30% (as per dm3 build) |
| MAR density per gene | 1 | 3 |

### 2.3 Motif analysis of intergenic sequences

The intergenic sequences from Chr2L,2R, 3L and 3R heterochromatic region were extracted using in-house Perl script and submitted for motif analysis using MEME [33]. The motif search was run with both default parameters (resulted in 50bp motif) and modified search for small motif of 15bp. The motif, thus, obtained was contributed by 53 out of 105 sequences submitted, 10 sequences had multiple occurrence of the motif. We queried the motifs against database of known motifs using TOMTOM.

### 2.4 Creation of Drosophila Het C-state

We have previously developed a visualization tool called C-State [27] to analyze multiple chromatin and expression datasets at once. This was customized such that it automatically loads the histone marks and expression data of *Drosophila* heterochromatic genes across various developmental stages <u>(Drosophila Het C-State)</u>. C-State fetches the relevant genomic information based on the input identifiers from its genome repository folder. Once genomic coordinates of all genes were retrieved, histone mark peaks mapping to the coordinates were extracted from the genome-wide data of each developmental stage, and transformed into coordinates relative to the TSS of the gene. C-State plots every gene was corrected for the genomic orientation, so that genes on both the positive and negative strands of the genome can be compared directly. Similarly, the expression level of each gene was retrieved from the genome-wide expression data for each developmental stage.

Average feature profiles are calculated using a constant-binning approach. First, each gene is divided into equal number of bins irrespective of its size and orientation, such that the TSS of all genes are represented by the same bin. Starting from the upstream most bin, the number of peaks mapping to each bin for all genes is calculated. Multiple peaks mapping to the same bin are counted only once, to avoid biases created by large bin sizes. Once the number of peaks for each bin is derived, they were plotted as line charts using a common Y-scale, to directly visualize the histone peak trends in a comparable manner.The “Files” interface was replaced with buttons that enable toggling between the display of data across all developmental. However the C-State json file is provided as Drosophila HetCState file.json for the ease of including new heterochromatic genes and ChIP datasets by the programmers. All the remaining options of C-State such as the filters, tables, plots and customization options are functional, and can be utilized by the user according to their convenience. The gene-epicard can be viewed at the Table summary tab.

## 3. Results

### 3.1 Epigenetic profile of heterochromatic genes across developmental stages is dynamic

Pericentromeric heterochromatin is reported to have several hundred genes [20]. We chose 60 representative heterochromatic genes that fall within autosomal chr2 and 3 pericentromeres as per Drosophila Heterochromatin Genome Project and their gene-structure; function and expression data was available. Some of these genes are studied extensively in the field of heterochromatic gene regulation-eg: *light, rolled, RpL(s), Nipped* etc. To investigate if these genes are subjected to dynamic epigenomic changes, we looked into the profile of six histone modifications across 12 well-established developmental stages: (E0-4, E4-8, E8-12, E12-16, E16-20, E20-24, L1, L2, L3, pupae, adult female and adult male) [34] using published datasets from modENCODE. Histone modification marks included are– two “inactive”: H3K9me3, H3K27me3 and four “active”: H3K4me1, H3K4me3, H3K9ac and H3K27ac, with corresponding developmental-stage specific ChIP seq/ ChIP-chip (**Table1**) and RNAseq datasets (refer to <u>Drosophila Het C-State</u>, for the trend of histone marks present on individual heterochromatic genes across developmental stages). Chr4 and X heterochromatic genes were excluded from the analysis as they were reported to have other specific marks like H4K16ac [35] for which datasets across all developmental stages were unavailable.

We analyzed the enrichments of six histone marks on the heterochromatic genes at their different levels of gene-expression. (**Fig1**) shows the average trend of their occurrence across the 12 developmental stages. The heterochromatin marker, H3K9me3, is present throughout the gene body with lesser enrichment in larval (L3) and adult stages (male and female). H3K4me3 was consistently present on most genes at their TSS except in L1. H3K4me1 is present mostly in embryo12-16 hours and rest of the stages show minimal enrichment at the TSS. H3K9and H3K27 acetylation peaks at the TSS with gradual tapering on the gene body, larval stage onwards consistent with the fact that most genes have low expression in larval and adult stages *(eg:CG17691, CG17683, uex, vtd, CG40006)*. [Supplementary Table 1: Stagewise expression data]. We found H3K27me3 was absent on most of the genes. Constitutively expressed heterochromatic genes like *(RpL5)* have less of H3K9me3, restricted mostly at the introns. H3K27me3 is present in the lowest / no expression stages of certain genes. The active marks H3K4me3 is present at the exons or throughout the gene body while H3K9/27Ac is also present at the gene body and the UTRs. For the heterochromatic genes with tissue/stage specific expression pattern (*Nipped-A*), during highest expression stage, active marks are present at the coding regions and repressive marks at the introns, while the entire gene gets marked with H3K9me3 marks during lower/no expression stages. This trend in occurrence of different histone marks is correlated to the expression pattern of these across development and was used to classify the genes in the following section

**Figure 1A:**
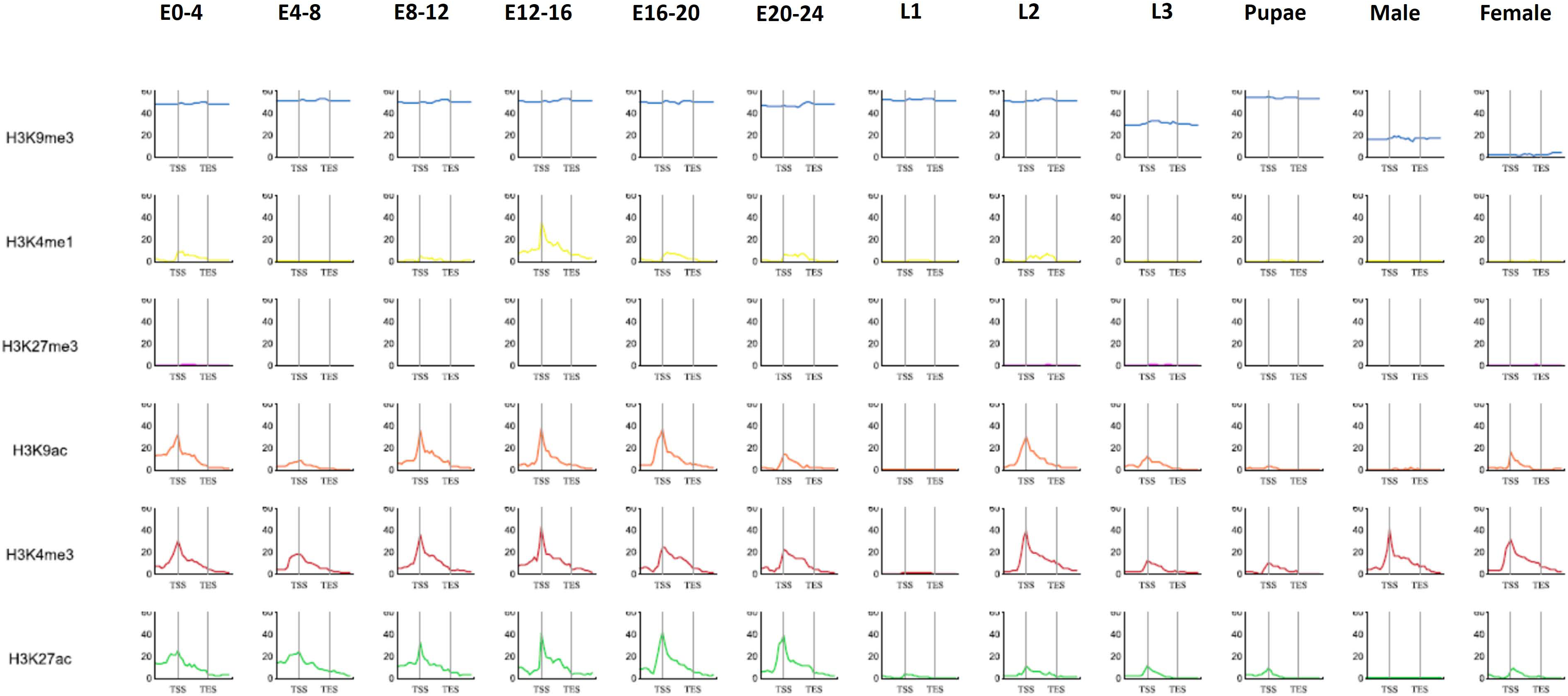
Average feature profile plot of histone modifications on heterochromatic genes across all developmental stages depicting the average trend of occurrence of six histone modifications over the entire gene body (TSS to TES marked by black vertical bars) for the 12 developmental stages. H3K9me3 remains consistently present on the gene body with significant lowering only in adult stages. H3K4me1 shows a peak mostly in the embryos12–16hrs whereas H3K4me3 has a constant peak at the TSS followed by gradual tapering at all stages except L1. H3K9ac and H3K27ac also shows a sharp peak at the TSS except for in the larval, pupal and adult male stages. H3K27me3 is absent on most of the heterochromatic genes

### 3.2 Classification of heterochromatic genes based on the dynamics of repressive histone marks

We were interested to understand the dynamics of histone modifications and correlate it with the expression pattern that might not be evident by looking at the cumulative trend across all stages. Thus, we divided the expression datasets into two stages: highest and lowest/no expression for each gene. **Fig2A** shows the average trend in the distribution of the histone marks on the heterochromatic genes at their highest and lowest stages of expression. The typical feature of the heterochromatic genes that sets it apart from the euchromatic genes is the presence of a combination of active and inactive histone modifications [Supplementary Figure 1]. We sought to group the genes based on the presence or absence of inactive H3K9me3 marks in the stages of highest and lowest expression. We grouped the genes into three classes as shown in **Table 2**. These results show that genes of each group depend on the inactive histone marks for their expression to different extents (**Fig2B)**. Group I-A genes have both inactive and active histone marks in stages of highest expression and only inactive in lowest stages while Group I-B have both kinds of histone marks in both highest and lowest expression. Genes in both these sub-classes are majorly (92%) constitutively expressing (**Table 2**) and encode proteins involved in metabolic and developmental pathways. Notable examples are ribosomal proteins *(RpL 5/15/38)*, kinases *(Stlk)* and proteins involved in signaling *(Gprk1)*, cell-divison *(Nipped-B, uex)* and transcriptional regulation *(Atf6, CG10395, Maf1)*. Group I also constitutes the largest class with 58% of the genes included in our study. Group II genes consisting of 10% of genes studied, showed enrichment of only active marks in highest and inactive marks in lowest stages of expression respectively. Group III genes had only inactive histone marks in the both the stages. Interestingly, almost 50% of the genes in this class had tissue-specific expression having role in metabolic processes and transferase activity implicated in wing and muscle development.

**Figure 2A:**
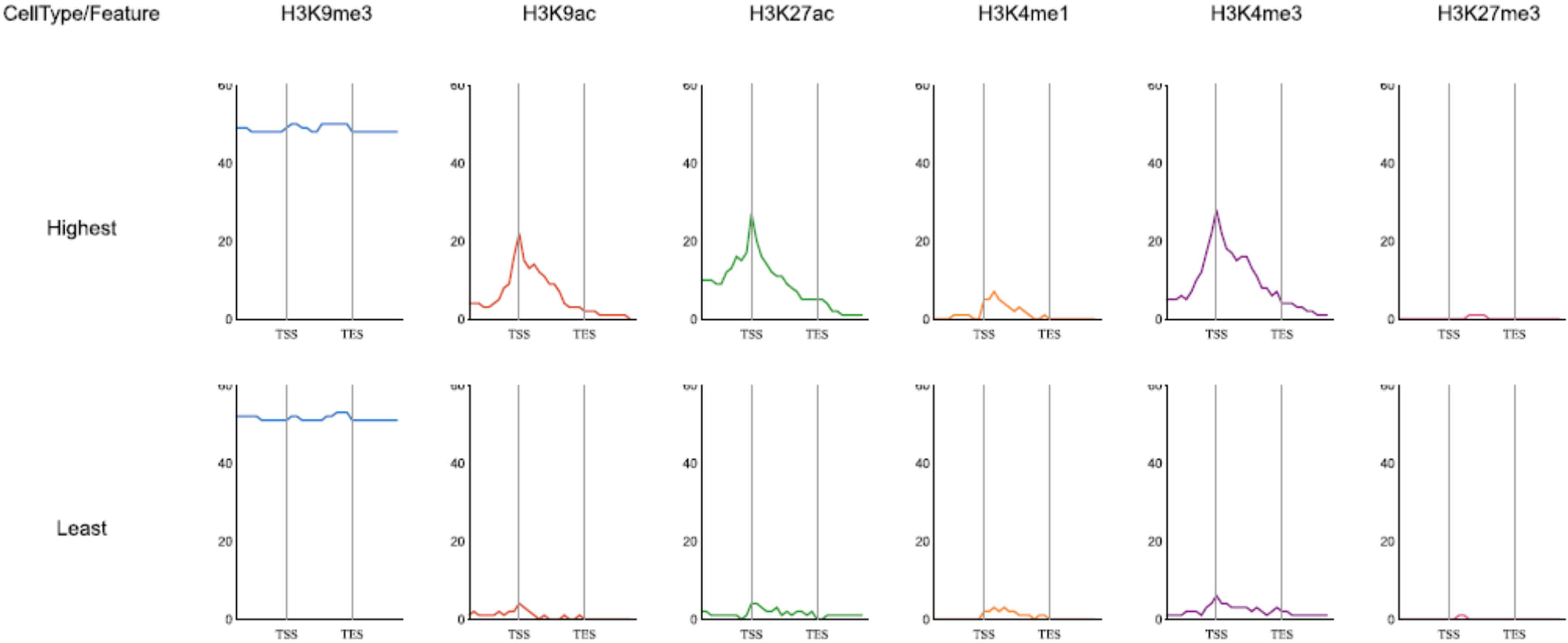
Average plot for the distribution of the histone marks in the highest and least stages of expression. The stages of highest expression are marked by the presence of H3K9me3 throughout the gene-body with sharp peak of other active marks like H3K9/27ac and H3K4me1/3 at the TSS. At the stages of lowest expression H3K9me3 prevails while the level of the active histone marks go down significantly. Since this is an average plot for all genes meaning it includes constitutively active genes so the enrichment levels of the active marks in the least expression stage do not go down to 0.

**Figure 2B:**
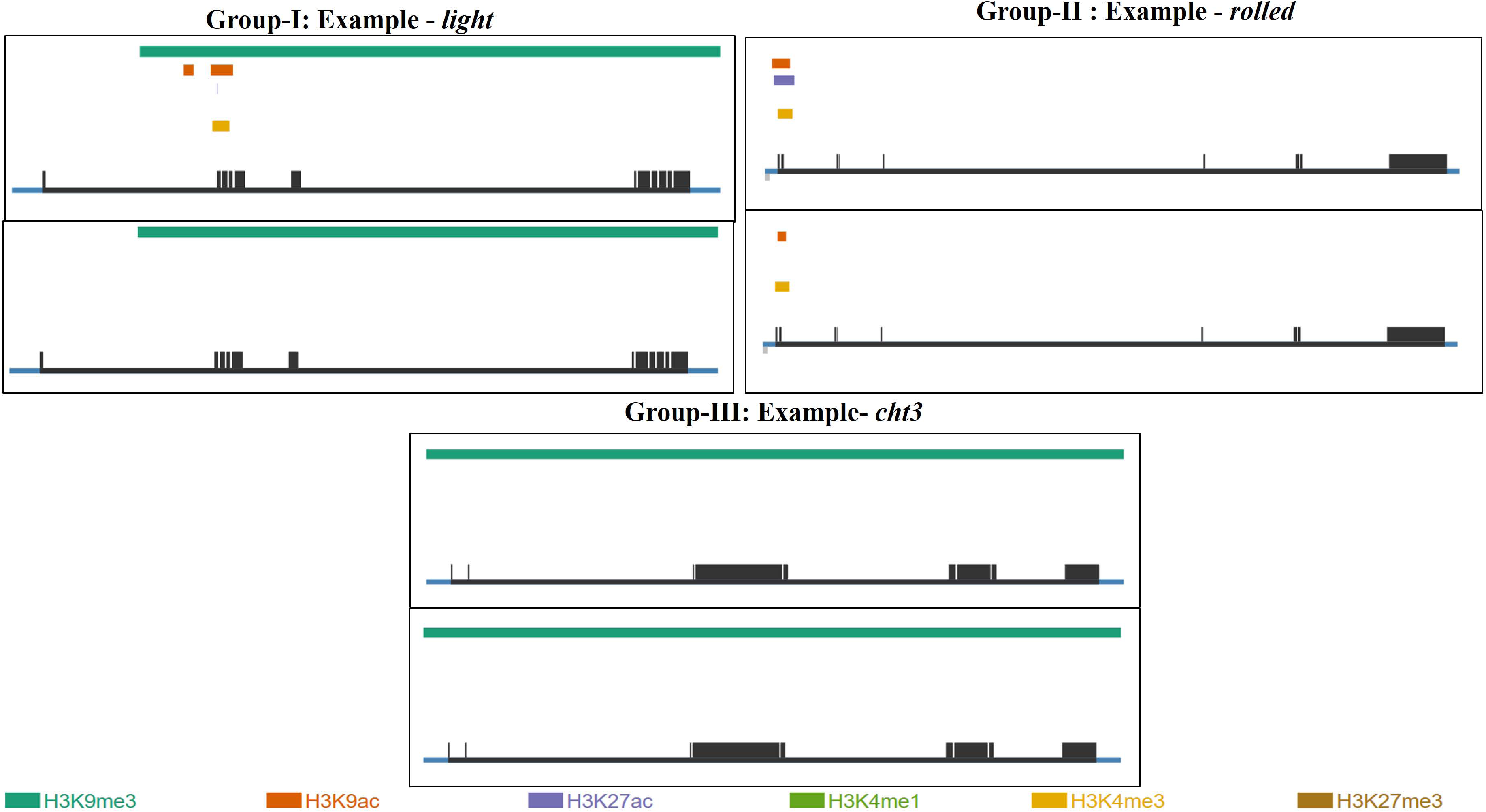
Representative example from each Group of genes at their stages of highest and least expression. Groupl: Example-light – active and inactive histone marks in stages of highest expression and only inactive mark of H3K9me3 in lowest expression stages; Group II: Example-*rolled* shows presence of active histone marks in both stages of highest and lowest expression; GroupIII:Example-*cht3* has H3K9me3 in highest and lowest stages of expression

### 3.3 Heterochromatic genes have preferential enrichment of Matrix Associated Regions in the pericentromeric intergenic regions

Regulatory DNA elements along with the histone modifications modulate the transcriptional status of genes. We hypothesized that MARs might regulate genome organization in pericentromere, looping out active genes from neighboring heterochromatin into separate chromatin domains. We have previously published the genome-wide sequencing of MARs in the *Drosophila* embryo (0–16hrs) and their features [36]. However, MARs mapping to the pericentromeric regions were not included. Hence, we were interested to determine the characteristics of those Het-MARs. We took up all the MARs that fall within the pericentromeric heterochromatin, including those that map to centromere as per the latest genome build -dm6 annotation [31]. 350 MARs were mapped to pericentromeric regions of the autosomal chromosomes and X chromosome. We compared the properties of MARs with those reported previously in our study for the euchromatic regions as shown in **Table3**. We found that the overall average size of Het MARs is 354 bp, almost half of the size in the euchromatic region. The distribution of the MARs with respect to the various genomic features (**Fig3A)** shows that most pericentromeric heterochromatin are intergenic. We also looked at the inter-Het MAR distance, the average being 45 Kb and the largest being close to few Mb. The MARs/genes value indicates that MARs can probably loop out few similarly expressing genes into separate domain. (**Fig3B**) shows the genomic map of the distribution of MARs on the chromosome arms along with the histone marks indicating that in certain cases MARs are present at the borders of distinct chromatin domains.

**Figure 3A:**
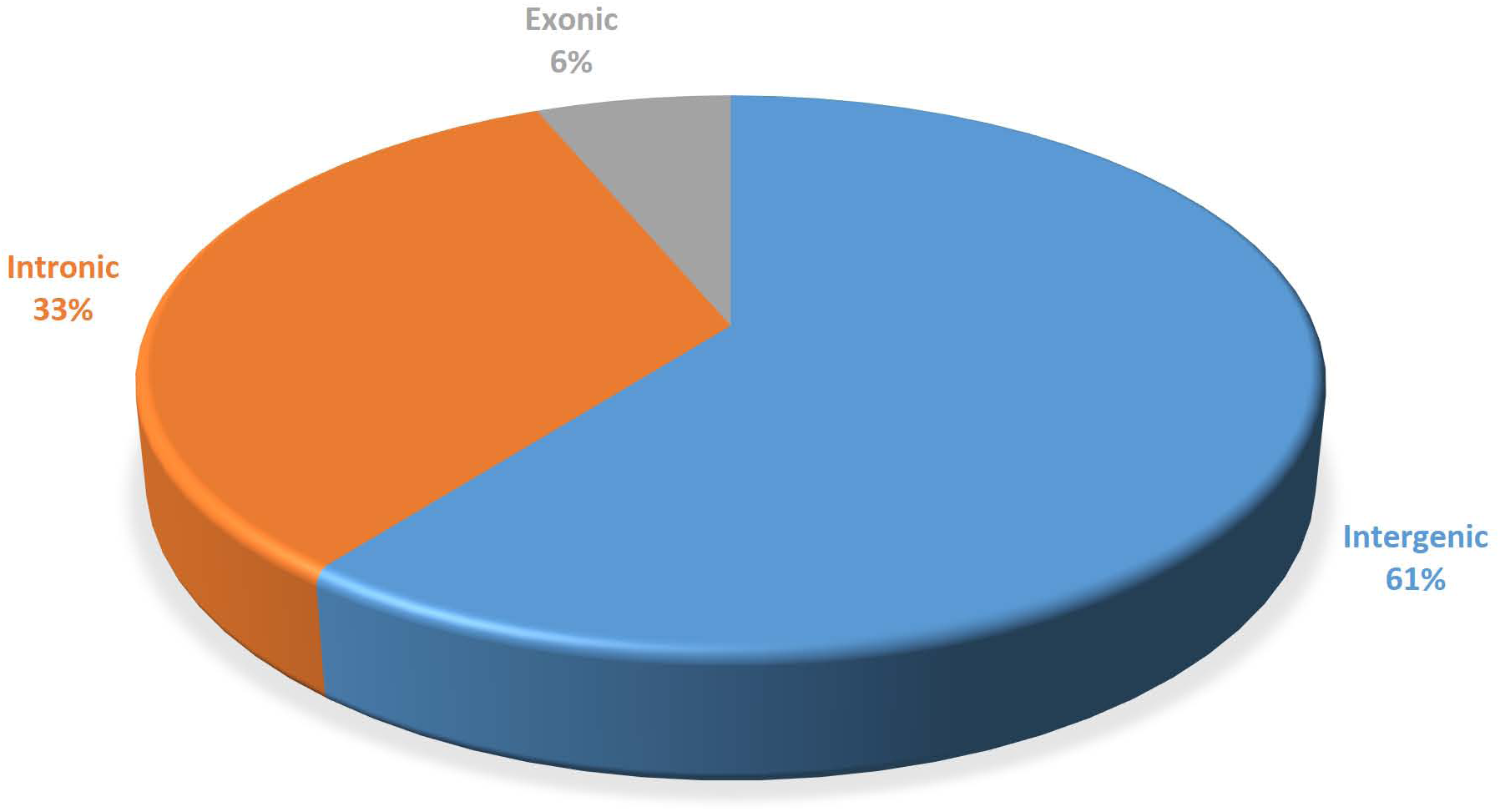
Distribution of HetMARs in pericentromeric heterochromatin in pericentromeric heterochromatin with respect to various genic features

**Figure 3B:**
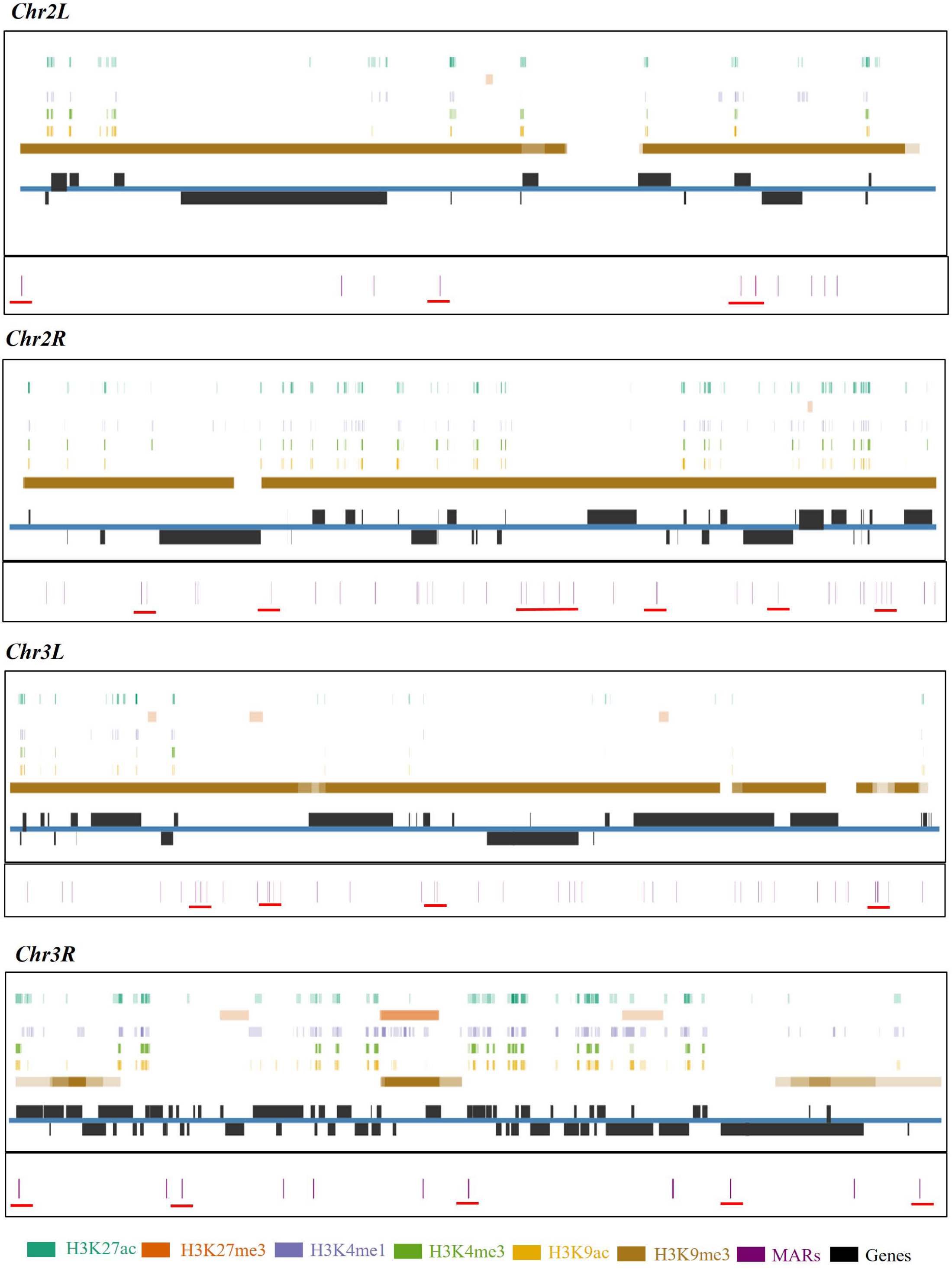
The distribution of Het MARs on 4 chromosomal pericentromeric heterochromatin regions where genes are shown in black and the 0–16 hrs embryo MARs in violet. Different histone modifications from 0–16hrs are shown in different colors. The higher the peak intensity of each histone mark across the (0-4h, 4-8h, 8-12h and 12-16h) 4 stages, less opaquer is their respective bars. MARs at the intergenic regions of chromatin domain borders are highlighted by red

### 3.4 Heterochromatic intergenic sequences have DNA motifs of transcription factors

To investigate whether any specific motifs are enriched at the pericentromeric heterochromatin, intergenic sequences of Chr 2 and 3, pericentromeric genes were extracted and used for motif analysis using MEME by the default parameters. We found a 15 bp motif to be highly enriched in the intergenic sequences with a dependable e value 1.9e – 423 and an occurrence in 70% sequences studied as seen in (**Fig4)**. The motif matched to known motifs of Zn finger DNA binding proteins - CG3065, Sp1 and CG12029. Of these hits, particularly interesting is the presence of motifs for binding of Sp1 a well-known transcription factor that is known to modulate epigenetic environment by interacting with several chromatin remodelers.

**Figure 4:**
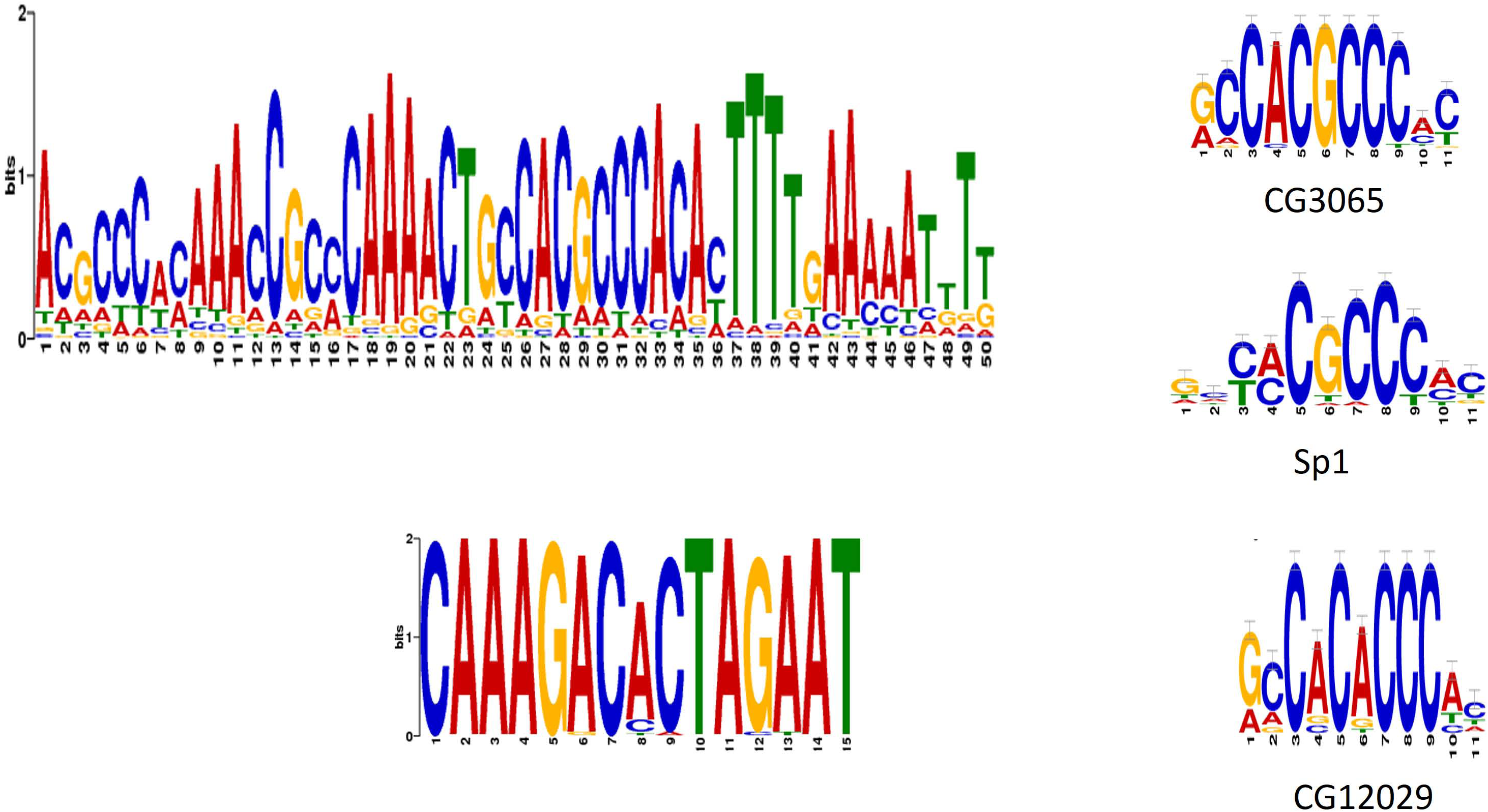
MEME analysis of intergenic sequences of the heterochromatic genes on Chr2 and 3. The MEME analysis was ran using default parameters that resulted in 50bp motif (above) and using modified parameters that gives a 15bp motif with a E value: 1.9e-432 (below). The smaller motif is overlapping with the 50bp motif. 15bp motif was queried against the database of known motifs using TOMTOM. Top hits were of CG3065, Sp1 and CG12029 - all Zn finger DNA binding domain containing transcription factors.

## Discussion

It is counterintuitive to have genes in the repressed chromatin environment that get expressed by adapting and utilizing the heterochromatic factors rather than being aversive to them. To understand this paradigm, we used Drosophila Het C-State to compare trends of histone modifications on heterochromatic genes across development. This also serves as the repository for information specifically related to Drosophila heterochromatic genes to which new coordinate based genomic datasets or heterochromatic genes can be added to cater to the community of researchers working on these genes.

Previous studies had shown that in normal embryos H3K9me2 is enriched throughout the gene body while in embryos with chromosomal rearrangement that created a new eu-het junction-H3K9me2 distribution is altered [37]. More recently, epigenetic landscape of the heterochromatic genes was investigated in S2, BG3 cells and embryos using ChIP-chip data of several histone marks. Collectively these studies showed that active heterochromatic genes have unique combinations-enrichment of active (H3K9/14ac and H3K4me2) marks with a dip in H3K9me3 at the TSS [37] and the presence of both active (H3K36me3) and inactive (H3K9me3) marks on the gene body, not seen in case of euchromatic genes [35]. Previous studies had used tiling array based ChIP data and focused on a single cell type or developmental stage. Therefore, how the histone marks change with respect to changing expression patterns across development was unexplored. Our results show that the distribution of H3K4me1/3, H3K9/27ac and H3K9me3 on the heterochromatic genes is different from euchromatic genes such that both these active marks and inactive mark of H3K9me3 occur together [Supplementary Fig 1]. This shows that there could be different combinations of active marks - H3K4me3-H3K9/27ac along with H3K9me3, which was not, reported earlier (**Fig1)**. We believe that the dip in H3K9me3 but presence of H3K9/27ac at the TSS is required to allow access of transcriptional machinery following which different combinations of active marks are put to mark the gene for expression. H3K4me1 promotes chromatin accessibility and is present (on average) at the TSS only in the highest expression stage. (**Fig2A)**. As the distinctive feature of heterochromatic genes, H3K9me3 is present in both the high and low expression stages. To probe further, as to how the heterochromatic mark: H3K9me3 regulates both activation and repression of heterochromatic genes we looked into the genes groupwise.

We grouped the genes based on the presence of heterochromatic mark during active expression (**Table 2)**. Group I genes include most of the constitutive genes that have active marks like H3K9/27ac at TSS and H3K4me3 at the gene body along with H3K9me3. In stages of lower expression, there is decrease in presence of active marks but H3K9me3 persists. Inactive marks on the 5’end of the gene are probably read differently than those on the gene body that was reported to occur in combination with other active marks [35,38]. The second category (Group II) of genes had only active marks in both highest and lowest expression stages. These genes are like euchromatic genes embedded in heterochromatin. We also found the well-known heterochromatic gene *-rolled* in this class. Probably, they need other heterochromatic factors like HP1, Su(var)3-9 for expression although H3K9me3 is not present on the gene body and thus the pattern of histone marks on them resembles those on euchromatic genes. Greil and colleagues had reported the binding of HP1a and Su(var)3-9 on *rolled* and *light* but upon knockdown of these two proteins in Kc167 cells they do not see change in expression level of these 2 genes [39]. Hence how HP1a and Su(var)3-9 controls heterochromatic genes expression needs further experimental dissections. Group III genes showed enrichment for only inactive marks in stages of active expression. To explain that only inactive mark-H3K9me3 is present on transcriptionally active group III genes, we consider two possibilities. First, in mammals H3K9me3 has also been reported to promotes transcriptional elongation in concert with HP1c [40]. In fly heterochromatic genes H3K36me3 (elongation mark) has been shown to be present along with H3K9me3 on the gene-body [35]. However, how it regulates expression is not understood. Second, the major heterochromatic factor HP1 that binds to H3K9me3, has been shown to be present on active pericentric genes [41]. HP1 is known to have gene activation effect [42] through its differential post-translational modifications [43] and interaction partners [44] including MES-4, a H3K36 methyl-transferases. Hence, the role of HP1a in regulation of Group III gene expression needs to be explored.

In summary, these three classes of genes are regulated to different extents by the heterochromatic factors or has combinations of other active histone modifications. Thus, resulting in a complex crosstalk of multiple pathways being involved in the context of heterochromatic gene expression. This highlights the need for meta-analysis of available genomic datasets to bring out the specialties of heterochromatic genes and test out hypotheses experimentally.

Among the several models proposed for the regulation of the heterochromatic genes, ‘Integration model’ postulates that the long-range interactions mediated by the heterochromatic proteins and sequences can explain context dependent regulation of heterochromatic genes [45]. It is known that the co-expression clusters are encompassed within topologically associated domains (TADs) that can be of 40–70kb in *Drosophila* [46]. Although based on the epigenetic landscape, it is believed that centromeric heterochromatic regions are folded into a compact inactive TAD, the detailed characterization of long-range interactions in pericentromeric associated domains as reported for mice [47], is lacking for *Drosophila*. MARs tether the genome to the nuclear matrix thus determining the folding of the genome into chromatin domains. MARs have been shown to promote long range enhancer-promoter interactions even in the repressive environment [48]. However, their role in the context of heterochromatic gene regulation is unexplored. In this backdrop, we looked into the distribution of MARs within the pericentromeric heterochromatin. Our analysis shows preferential enrichment of MARs in intergenic regions with the inter Het MARs distance in the range of average *Drosophila* TAD size. MARs have shown to modulate chromatin accessibility [49] and in many cases associate with actively transcribing genes [50]. More recently MARs were associated with active transcription [50,51] and that inter-MAR looping contributes to transcriptionally active DNA looping [52]. Thus, it can play a role in mediating long-range interaction to keep heterochromatic genes active. We propose that MARs could be defining the borders in pericentromeric heterochromatin that confines long-range interactions between similarly expressing heterochromatic genes into separate chromatin domains. However, this observation needs further experimental validations.

Transcription factor binding sites (TFBS) and transcription factor dosage has also been shown to remodel chromatin landscape and impact transcriptional status of the genes [53]. The presence of motif recognized by a transcription factor like Sp1 points to the significance of non-coding intergenic sequences in shaping the chromatin environment. Sp1 through to its DNA binding domain interacts with acetyltransferase domain of co-activator p300 to increase the gene expression [27]. In addition, Sp1 binding is involved in de-acetylation and cause repression of several genes [28]. The presence of Sp1 binding motifs at the intergenic regions can be speculated to modulate the chromatin environment. However, experimental validations of this hypothesis must ascertain how the combination of non-coding sequences along with *trans* factors regulate the epigenetic landscape of centromeric heterochromatin.

In conclusion, this is the first report of the dynamic epigenetic landscape of *D melanogaster* heterochromatic genes during development. We present Drosophila Het C-State as platform for comprehensive bioinformatics analyses using the publicly available genomics datasets and bring out the peculiarities of heterochromatic genes that might be involved in their regulation. Heterochromatic genes are also known in mammals and many of them are expressed during early development [4,54] and disease conditions like cancer [3,55]. We believe that despite the inherent challenges of studying heterochromatic sequences due to high repeat content and compaction, more experimental evidences are required to explain these observations. Such studies will lead us to understand the dynamics of heterochromatic gene regulation and shed more light into the dark matter of the genome.

## Acknowledgements

We thank Rashmi Upadhyay Pathak and A Srinivasan for their help with the MAR dataset. We thank Surabhi Srivastav for critically reading the manuscript. PS thanks University Grants Commision (UGC), India for the doctoral fellowship. DTS and RKM acknowledges the financial support of the Council of Scientific and Industrial Research (CSIR), India.

---

**Supplementary Figure 1: Distinction between euchromatic and heterochromatic epigenomic landscape-** shows that the euchromatic regions there is presence of only active marks viz, H3K27ac, H3K9ac and H3K4me1/3. At some places even facultative repressive mark H3K27me3 is present. However, on the right side of the dotted line demarcates a heterochromatic region where H3K9me3 is present along with the other active marks mostly at the TSS – an unique feature of the epigenomic landscape of heterochromatic genes. Notably, the gene density is lower in heterochromatin as compared to heterochromatin

## References

[1] C.D. Smith, S. Shu, C.J. Mungall, G.H. Karpen, The Release 5.1 annotation of Drosophila melanogaster heterochromatin., Science. 316 (2007) 1586–1591. doi:10.1126/science.1139815.

[2] M.E. Brun, M. Ruault, M. Ventura, G. Roizès, A. De Sario, Juxtacentromeric region of human chromosome 21: A boundary between centromeric heterochromatin and euchromatic chromosome arms, Gene. 312 (2003) 41–50. doi:10.1016/S0378-1119(03)00530-4.

[3] M.E. Brun, E. Lana, I. Rivals, G. Lefranc, P. Sarda, M. Claustres, A. Mégarbané, A. de Sario, Heterochromatic genes undergo epigenetic changes and escape silencing in immunodeficiency, centromeric instability, facial anomalies (ICF) syndrome, PLoS One. 6 (2011) 1–8. doi:10.1371/journal.pone.0019464.

[4] M.C. Beckers, J. Gabriëls, S. Van Der Maarel, A. De Vriese, R.R. Frants, D. Collen, A. Belayew, Active genes in junk DNA? Characterization of DUX genes embedded within 3.3 kb repeated elements, Gene. 264 (2001) 51–57. doi:10.1016/S0378-1119(00)00602-8.

[5] T.N. Le, Y. Miyazaki, S. Takuno, H. Saze, Epigenetic regulation of intragenic transposable elements impacts gene transcription in Arabidopsis thaliana, Nucleic Acids Res. 43 (2015) 3911–3921. doi:10.1093/nar/gkv258.

[6] P. Dimitri, N. Corradini, F. Rossi, F. Vernì, The paradox of functional heterochromatin, BioEssays. 27 (2005) 29–41. doi:10.1002/bies.20158.

[7] P. Dimitri, R. Caizzi, E. Giordano, M. Carmela Accardo, G. Lattanzi, G. Biamonti, Constitutive heterochromatin: A surprising variety of expressed sequences, Chromosoma. 118 (2009) 419–435. doi:10.1007/s00412-009-0211-y.

[8] A.J. Hilliker, Genetic analysis of the centromeric heterochromatin of chromosome 2 of Drosophila melanogaster: deficiency mapping of EMS induced lethal complementation groups, Genetics. 83 (1976) 765–782.

[9] D.E. Koryakov, I.F. Zhimulev, P. Dimitri, Cytogenetic analysis of the third chromosome heterochromatin of Drosophila melanogaster, Genetics. 160 (2002) 509–517.

[10] N. Corradini, F. Rossi, E. Giordano, R. Caizzi, F. Verní, P. Dimitri, Drosophila melanogaster as a model for studying protein-encoding genes that are resident in constitutive heterochromatin., Heredity (Edinb). 98 (2007) 3–12. doi:10.1038/sj.hdy.6800877.

[11] P. Dimitri, N. Corradini, F. Rossi, F. Vernì, G. Cenci, G. Belloni, I.F. Zhimulev, D.E. Koryakov, Vital genes in the heterochromatin of chromosomes 2 and 3 of Drosophila melanogaster, Genetica. 117 (2003) 209–215. doi:10.1023/A:1022996112742.

[12] S. Schulze, D. a R. Sinclair, E. Silva, K. a. Fitzpatrick, M. Singh, V.K. Lloyd, K. a. Morin, J. Kim, D.G. Holm, J. a. Kennison, B.M. Honda, Essential genes in proximal 3L heterochromatin of Drosophila melanogaster, Mol. Gen. Genet. 264 (2001) 782–789. doi:10.1007/s004380000367.

[13] K. a Fitzpatrick, D. a Sinclair, S.R. Schulze, M. Syrzycka, B.M. Honda, A genetic and molecular profile of third chromosome centric heterochromatin in Drosophila melanogaster., Genome. 48 (2005) 571–584. doi:10.1139/g05-025.

[14] S.R. Schulze, D. a R. Sinclair, K. a. Fitzpatrick, B.M. Honda, A genetic and molecular characterization of two proximal heterochromatic genes on chromosome 3 of Drosophila melanogaster, Genetics. 169 (2005) 2165–2177. doi:10.1534/genetics.103.023341.

[15] S.R. Schulze, B.F. McAllister, D. a R. Sinclair, K. a. Fitzpatrick, M. Marchetti, S. Pimpinelli, B.M. Honda, Heterochromatic genes in Drosophila: A comparative analysis of two genes, Genetics. 173 (2006) 1433–1445. doi:10.1534/genetics.106.056069.

[16] A.B. Coulthard, C. Alm, I. Cealiac, D. a. Sinclair, B.M. Honda, F. Rossi, P. Dimitri, A.J. Hilliker, Essential loci in centromeric heterochromatin of Drosophila melanogaster. I: The right arm of chromosome 2, Genetics. 185 (2010) 479–495. doi:10.1534/genetics.110.117259.

[17] D.A.R. Sinclair, S. Schulze, E. Silva, K.A. Fitzpatrick, B.M. Honda, Essential genes in autosomal heterochromatin of Drosophila melanogaster, Genetica. 109 (2001) 9–18. doi:10.1023/A:1026500620158.

[18] S. Pimpinelli, S. Bonaccorsi, M. Gatti, L. Sandler, The peculiar genetic organization of Drosophila heterochromatin, Trends Genet. 2 (1986) 17–20. doi:10.1016/0168-9525(86)90163-0.

[19] F.L. Sun, M.H. Cuaycong, C.A. Craig, L.L. Wallrath, J. Locke, S.C. Elgin, The fourth chromosome of Drosophila melanogaster: interspersed euchromatic and heterochromatic domains., Proc. Natl. Acad. Sci. U. S. A. 97 (2000) 5340–5. doi:10.1073/pnas.090530797.

[20] R. a Hoskins, J.W. Carlson, C. Kennedy, D. Acevedo, M. Evans-Holm, E. Frise, K.H. Wan, S. Park, M. Mendez-Lago, F. Rossi, A. Villasante, P. Dimitri, G.H. Karpen, S.E. Celniker, Sequence finishing and mapping of Drosophila melanogaster heterochromatin., Science. 316 (2007) 1625–1628. doi:10.1126/science.1139816.

[21] J.C. Yasuhara, C.H. DeCrease, B.T. Wakimoto, Evolution of heterochromatic genes of Drosophila., Proc. Natl. Acad. Sci. U. S. A. 102 (2005) 10958–10963. doi:10.1073/pnas.0503424102.

[22] D.F. Eberl, B.J. Duyf, A.J. Hilliker2, The Role of Heterochromatin in the Expression of a Heterochromatic Gene, the rolled Locus of Drosophila melanogaster, (n.d.).

[23] B.T. Wakimoto, M.G. Hearn, The effects of chromosome rearrangements on the expression of heterochromatic genes in chromosome 2L of Drosophila melanogaster., Genetics. 125 (1990) 141–54. http://www.pubmedcentral.nih.gov/articlerender.fcgi?artid=1203996&tool=pmcentrez&rendertype=abstract.

[24] B.Y. Lu, P.C. Emtage, B.J. Duyf, a J. Hilliker, J.C. Eissenberg, Heterochromatin protein 1 is required for the normal expression of two heterochromatin genes in Drosophila., Genetics. 155 (2000) 699–708.

[25] A.Y. Hessler, V-Type Position Effects At the Light Locus in Drosophila Melanogaster, (1957).

[26] N.J. Clegg, B.M. Honda, I.P. Whitehead, T. a Grigliatti, B. Wakimoto, H.W. Brock, V.K. Lloyd, D. a Sinclair, Suppressors of position-effect variegation in Drosophila melanogaster affect expression of the heterochromatic gene light in the absence of a chromosome rearrangement., Genome. 41 (1998) 495–503.

[27] D.T. Sowpati, S. Srivastava, J. Dhawan, R.K. Mishra, C-State: An interactive web app for simultaneous multi-gene visualization and comparative epigenetic pattern search, BMC Bioinformatics. (2017). doi:10.1186/s12859-017-1786-6.

[28] B.R. Graveley, A.N. Brooks, J.W. Carlson, M.O. Duff, J.M. Landolin, L. Yang, C.G. Artieri, M.J. van Baren, N. Boley, B.W. Booth, J.B. Brown, L. Cherbas, C.A. Davis, A. Dobin, R. Li, W. Lin, J.H. Malone, N.R. Mattiuzzo, D. Miller, D. Sturgill, B.B. Tuch, C. Zaleski, D. Zhang, M. Blanchette, S. Dudoit, B. Eads, R.E. Green, A. Hammonds, L. Jiang, P. Kapranov, L. Langton, N. Perrimon, J.E. Sandler, K.H. Wan, A. Willingham, Y. Zhang, Y. Zou, J. Andrews, P.J. Bickel, S.E. Brenner, M.R. Brent, P. Cherbas, T.R. Gingeras, R.A. Hoskins, T.C. Kaufman, B. Oliver, S.E. Celniker, The developmental transcriptome of Drosophila melanogaster, Nature. 471 (2011) 473–479. doi:10.1038/nature09715.

[29] Y. Hu, A. Comjean, N. Perrimon, S.E. Mohr, The Drosophila Gene Expression Tool (DGET) for expression analyses, BMC Bioinformatics. 18 (2017) 98. doi:10.1186/s12859-017-1509-z.

[30] S. Roy, J. Ernst, P. V Kharchenko, P. Kheradpour, N. Negre, M.L. Eaton, J.M. Landolin, C. a Bristow, L. Ma, M.F. Lin, S. Washietl, B.I. Arshinoff, F. Ay, P.E. Meyer, N. Robine, N.L. Washington, L. Di Stefano, E. Berezikov, C.D. Brown, R. Candeias, J.W. Carlson, A. Carr, I. Jungreis, D. Marbach, R. Sealfon, M.Y. Tolstorukov, S. Will, A. a Alekseyenko, C. Artieri, B.W. Booth, A.N. Brooks, Q. Dai, C. a Davis, M.O. Duff, X. Feng, A. a Gorchakov, T. Gu, J.G. Henikoff, P. Kapranov, R. Li, H.K. MacAlpine, J. Malone, A. Minoda, J. Nordman, K. Okamura, M. Perry, S.K. Powell, N.C. Riddle, A. Sakai, A. Samsonova, J.E. Sandler, Y.B. Schwartz, N. Sher, R. Spokony, D. Sturgill, M. van Baren, K.H. Wan, L. Yang, C. Yu, E. Feingold, P. Good, M. Guyer, R. Lowdon, K. Ahmad, J. Andrews, B. Berger, S.E. Brenner, M.R. Brent, L. Cherbas, S.C.R. Elgin, T.R. Gingeras, R. Grossman, R. a Hoskins, T.C. Kaufman, W. Kent, M.I. Kuroda, T. Orr-Weaver, N. Perrimon, V. Pirrotta, J.W. Posakony, B. Ren, S. Russell, P. Cherbas, B.R. Graveley, S. Lewis, G. Micklem, B. Oliver, P.J. Park, S.E. Celniker, S. Henikoff, G.H. Karpen, E.C. Lai, D.M. MacAlpine, L.D. Stein, K.P. White, M. Kellis, Identification of functional elements and regulatory circuits by Drosophila modENCODE., Science. 330 (2010) 1787–1797. doi:10.1126/science.1198374.

[31] R.A. Hoskins, J.W. Carlson, K.H. Wan, S. Park, I. Mendez, S.E. Galle, B.W. Booth, B.D. Pfeiffer, R.A. George, R. Svirskas, M. Krzywinski, J. Schein, M.C. Accardo, E. Damia, G. Messina, M. Méndez-Lago, B. De Pablos, O. V. Demakova, E.N. Andreyeva, L. V. Boldyreva, M. Marra, A.B. Carvalho, P. Dimitri, A. Villasante, I.F. Zhimulev, G.M. Rubin, G.H. Karpen, S.E. Celniker, The Release 6 reference sequence of the Drosophila melanogaster genome, Genome Res. 25 (2015) 445–458. doi:10.1101/gr.185579.114.

[32] H. Attrill, K. Falls, J.L. Goodman, G.H. Millburn, G. Antonazzo, A.J. Rey, S.J. Marygold, Flybase: Establishing a gene group resource for Drosophila melanogaster, Nucleic Acids Res. 44 (2016) D786–D792. doi:10.1093/nar/gkv1046.

[33] T.L. Bailey, N. Williams, C. Misleh, W.W. Li, MEME: discovering and analysing DNA and protein sequence motifs, Nucleic Acids Res. 34 (2006) W369–373. doi:10.1093/nar/gkl198.

[34] N. Nègre, C.D. Brown, L. Ma, C.A. Bristow, S.W. Miller, U. Wagner, P. Kheradpour, M.L. Eaton, P. Loriaux, R. Sealfon, Z. Li, H. Ishii, R.F. Spokony, J. Chen, L. Hwang, C. Cheng, R.P. Auburn, M.B. Davis, M. Domanus, P.K. Shah, C. a Morrison, J. Zieba, S. Suchy, L. Senderowicz, A. Victorsen, N. a Bild, a J. Grundstad, D. Hanley, D.M. MacAlpine, M. Mannervik, K. Venken, H. Bellen, R. White, M. Gerstein, S. Russell, R.L. Grossman, B. Ren, J.W. Posakony, M. Kellis, K.P. White, A cis-regulatory map of the Drosophila genome., Nature. 471 (2011) 527–531. doi:10.1038/nature09990.

[35] N.C. Riddle, A. Minoda, P. V. Kharchenko, A. a. Alekseyenko, Y.B. Schwartz, M.Y. Tolstorukov, A. a. Gorchakov, J.D. Jaffe, C. Kennedy, D. Linder-Basso, S.E. Peach, G. Shanower, H. Zheng, M.I. Kuroda, V. Pirrotta, P.J. Park, S.C.R. Elgin, G.H. Karpen, Plasticity in patterns of histone modifications and chromosomal proteins in Drosophila heterochromatin, Genome Res. 21 (2011) 147–163. doi:10.1101/gr.110098.110.

[36] R.U. Pathak, A. Srinivasan, R.K. Mishra, Genome-wide mapping of matrix attachment regions in Drosophila melanogaster., BMC Genomics. 15 (2014) 1022. doi: 10.1186/1471-2164-15-1022.

[37] J.C. Yasuhara, B.T. Wakimoto, Molecular landscape of modified histones in Drosophila heterochromatic genes and euchromatin-heterochromatin transition zones, PLoS Genet. 4 (2008) 0159–0172. doi:10.1371/journal.pgen.0040016.

[38] S. Chantalat, A. Depaux, P. Héry, S. Barral, J.Y. Thuret, S. Dimitrov, M. Gérard, Histone H3 trimethylation at lysine 36 is associated with constitutive and facultative heterochromatin, Genome Res. 21 (2011) 1426–1437. doi:10.1101/gr.118091.110.

[39] F. Greil, I. Van Der Kraan, J. Delrow, J.F. Smothers, E. De Wit, H.J. Bussemaker, R. Van Driel, S. Henikoff, B. Van Steensel, Distinct HP1 and Su(var)3-9 complexes bind to sets of developmentally coexpressed genes depending on chromosomal location, Genes Dev. 17 (2003) 2825–2838. doi:10.1101/gad.281503.

[40] C.R. Vakoc, S. a Mandat, B. a Olenchock, G. a Blobel, Histone H3 lysine 9 methylation and HP1gamma are associated with transcription elongation through mammalian chromatin., Mol. Cell. 19 (2005) 381–91. doi:10.1016/j.molcel.2005.06.011.

[41] E. De Wit, F. Greil, B. Van Steensel, High-resolution mapping reveals links of HP1 with active and inactive chromatin components, PLoS Genet. 3 (2007) 0346–0357. doi:10.1371/journal.pgen.0030038.

[42] J.C. Eissenberg, S.C.R. Elgin, HP1a: a structural chromosomal protein regulating transcription., Trends Genet. 30 (2014) 103–10. doi:10.1016/j.tig.2014.01.002.

[43] G. LeRoy, J.T. Weston, B.M. Zee, N.L. Young, M.D. Plazas-Mayorca, B.A. Garcia, Heterochromatin protein 1 is extensively decorated with histone code-like post-translational modifications., Mol. Cell. Proteomics. 8 (2009) 2432–42. doi:10.1074/mcp.M900160-MCP200.

[44] A.A. Alekseyenko, A.A. Gorchakov, B.M. Zee, S.M. Fuchs, P. V. Kharchenko, M.I. Kuroda, Heterochromatin-associated interactions of Drosophila HP1a with dADD1, HIPP1, and repetitive RNAs, Genes Dev. 28 (2014) 1445–1460. doi:10.1101/gad.241950.114.

[45] J.C. Yasuhara, B.T. Wakimoto, Oxymoron no more: the expanding world of heterochromatic genes, Trends Genet. 22 (2006) 330–338. doi:10.1016/j.tig.2006.04.008.

[46] T. Sexton, E. Yaffe, E. Kenigsberg, F. Bantignies, B. Leblanc, M. Hoichman, H. Parrinello, A. Tanay, G. Cavalli, Three-dimensional folding and functional organization principles of the Drosophila genome, Cell. 148 (2012) 458–472. doi:10.1016/j.cell.2012.01.010.

[47] P.J. Wijchers, G. Geeven, M. Eyres, A.J. Bergsma, M. Janssen, M. Verstegen, Y. Zhu, Y. Schell, C. Vermeulen, E. De Wit, Characterization and dynamics of pericentromere-associated domains in mice, (2015) 958–969. doi:10.1101/gr.186643.114.Freely.

[48] W.C. Forrester, L. a Fernández, R. Grosschedl, L. a Ferna, repression of long-range enhancer – promoter interactions Nuclear matrix attachment regions antagonize methylation-dependent repression of long-range enhancer – promoter interactions, (1999) 3003–3014.

[49] T. Jenuwein, W.C. Forrester, L.A. Fernández-Herrero, G. Laible, M. Dull, R. Grosschedl, Extension of chromatin accessibility by nuclear matrix attachment regions, Nature. 385 (1997) 269–272. doi:10.1038/385269a0.

[50] M. a. Keaton, C.M. Taylor, R.M. Layer, A. Dutta, Nuclear scaffold attachment sites within ENCODE regions associate with actively transcribed genes, PLoS One. 6 (2011). doi:10.1371/journal.pone.0017912.

[51] A.K. Linnemann, A.E. Platts, S. a. Krawetz, Differential nuclear scaffold/matrix attachment marks expressed genes, Hum. Mol. Genet. 18 (2009) 645–654. doi:10.1093/hmg/ddn394.

[52] L. Wang, L.-J. Di, X. Lv, W. Zheng, Z. Xue, Z.-C. Guo, D.-P. Liu, C.-C. Liang, Inter-MAR association contributes to transcriptionally active looping events in human beta-globin gene cluster., PLoS One. 4 (2009) e4629. doi:10.1371/journal.pone.0004629.

[53] M. Lundgren, C.M. Chow, P. Sabbattini, A. Georgiou, S. Minaee, N. Dillon, Transcription factor dosage affects changes in higher order chromatin structure associated with activation of a heterochromatic gene, Cell. 103 (2000) 733–743. doi:10.1016/S0092-8674(00)00177-X.

[54] A. De Iaco, E. Planet, A. Coluccio, S. Verp, J. Duc, D. Trono, DUX-family transcription factors regulate zygotic genome activation in placental mammals, Nat. Genet. (2017). doi:10.1038/ng.3858.

[55] C. Grunau, J. Buard, M.E. Brun, A. De Sario, Mapping of the juxtacentromeric heterochromatin-euchromatin frontier of human chromosome 21, Genome Res. 16 (2006) 1198–1207. doi:10.1101/gr.5440306.

